# Re-emergence of Severe Acute Diarrhea Syndrome Coronavirus (SADS-CoV) in Guangxi, China, 2021

**DOI:** 10.1101/2022.07.11.499512

**Authors:** Yan-kuo Sun, Jia-bao Xing, Zhi-ying Xu, Han Gao, Si-jia Xu, Jing Liu, Di-hua Zhu, Yi-fan Guo, Bin-shuo Yang, Xiong-nan Chen, Ze-zhong Zheng, Heng Wang, Lang Gong, Edward C. Holmes, Gui-hong Zhang

**Author notes:** These authors contributed equally to the study (Yan-kuo Sun, Jia-bao Xing). **Correspondence:** Dr. Gui-hong Zhang, Dr. Edward C Holmes,.

## Abstract

Severe acute diarrhea syndrome coronavirus (SADS-CoV) has had a major impact on the swine industry in China, but has not been detected since 2019. Using real-time qPCR and metagenomic surveillance we identified SADS-CoV in a pig farm experiencing diarrheal disease. Genomic analysis supported the undetected circulation of SADS-CoV since 2019.

## Introduction

The ongoing COVID-19 pandemic has exposed major gaps in our knowledge of the diversity, evolution, ecology, and host range of coronaviruses (1–3). Concurrently, recent research directed toward coronaviruses has been instrumental in revealing the importance of bats as reservoirs for a diverse array of viruses, with occasional cross-species transmission to other hosts (4–7).

Severe acute diarrhea syndrome coronavirus (SADS-CoV), an emerging virus responsible for severe, acute diarrhea and dehydration in piglets, is one of the most important bat-derived alphacoronaviruses (8, 9). It was first detected in Guangdong province, China, in 2017, and then spread to neighboring Fujian and Jiangxi provinces, resulting in substantial economic damage to the pig industry (10). Retrospective analysis showed that its epidemiological history could be traced back to August 2016 (11). Intriguingly, following these outbreaks, the prevalence of SADS-CoV seemingly experienced a cliff-like descent, with no reports of the virus since 2019 (12). To determine whether SADS-CoV has remained undetected in Chinese swine populations we launched a surveillance project focusing on pig farms experiencing large-scale outbreaks of diarrheal disease. From this, we report the first re-appearance of SADS-CoV since 2019.

## The Study

From October 2019 to February 2021, we performed routine surveillance for SADS-CoV in China, sampling from the pig farms experiencing major outbreaks of diarrheal disease. In total, we collected 1365 diarrheal feces and intestinal samples from 23 provinces, five autonomous regions, and four municipalities, performing real-time quantitative reverse transcription PCR (real-time qPCR) to detect SADS-CoV No positive cases were detected. However, in May 2021, a fatal swine diarrhea disease outbreak was observed in an intensive scale pig farm in Guangxi province, with more than 3000 deaths in piglets. The clinical symptoms of the infected animals included severe and acute diarrhea, vomiting, and weight loss, with a fatality rate up to 100%. Using real-time qPCR on mixed samples in the farm we subsequently detected the aetiological agents for porcine epidemic diarrhea virus (PEDV), porcine deltacoronavirus (PDCoV), transmissible gastroenteritis virus (TGEV), as well as a single case of SADS-CoV

In the case of the SADS-CoV positive case, the cycle threshold (CT) was 23.18, indicative of a high viral load, enabling us to obtain the complete viral genome by metagenomic sequencing. Accordingly, following quality control and *de novo* assembly, we obtained the genome of a SADS-CoV that we have termed SADS-CoV/Guangxi/2021 (GeneBank Accession number. ON911569). This genomic sequence exhibited 98.4%-98.8% identity to the SADS-CoVs reported previously, and 87.4%-93.9% similarity with the related bat-borne HKU2 coronavirus. The virus exhibited high sequence similarity with previously sampled SADS-CoV across the entire genome, with the exception of the more diverse structural protein regions.

To determine the epidemiological origins of CoV/Guangxi/2021 we performed a phylogenetic analysis. First, to place the evolution of SADS-CoV in a broader context, we estimated a maximum likelihood (ML) phylogeny of the genus *Alphacoronavirus* as a whole (Figure 1A). As expected, this revealed that SADS-CoV fell within a diversity of coronaviruses sampled from *Rhinolophus* bats, including HKU2, and indicative of a cross-species transmission event. A more detailed complete genome phylogenetic analysis of the SADS-CoV cluster revealed that SADS-CoV/Guangxi/2021 fell as a distinct lineage within this group (Figure 1B). However, in a phylogenetic analysis of the most variable spike (S) gene, SADS-CoV/Guangxi/2021 clustered with virus sequences sampled from Jiangxi province, although characterized by a relatively long branch length (Appendix). This phylogenetic pattern suggests that SADS-CoV/Guangxi/2021 was derived from the ongoing transmission and evolution of SADS-CoV in China, rather than a separate cross-species transmission from bats, with the long branch length indicative of unsampled evolution since 2019 as well as the possible impact of recombination (see below). In addition, that SADS-CoV/Guangxi/2021 grouped with viruses from Jiangxi province in the S gene is indicative of inter-provincial movement, although the small available sample size prevents the exact identification of geographic origins.

**Figure 1.**
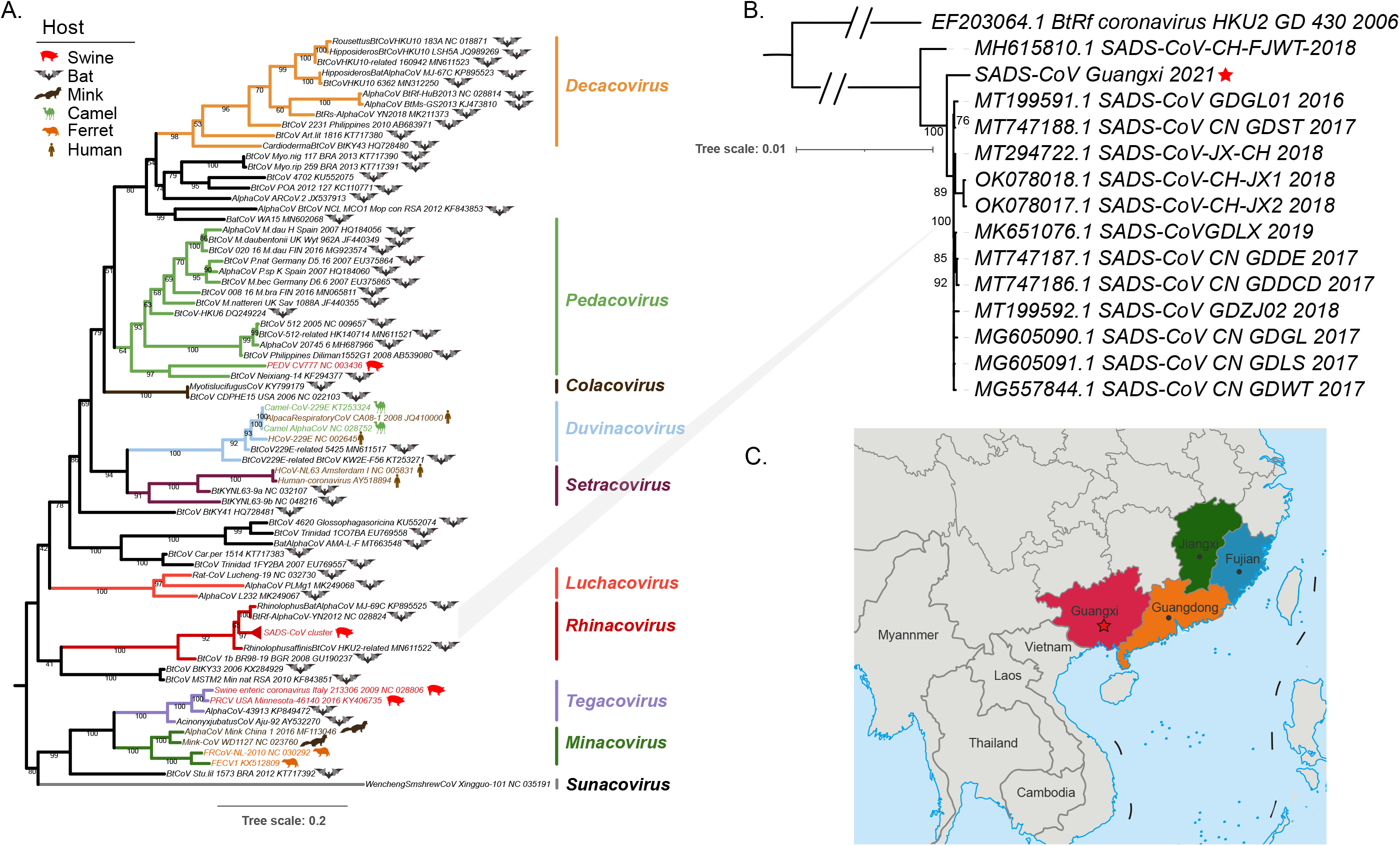
Evolutionary history of SADS-CoV. (A) Maximum likelihood phylogeny (IQ-TREE) of alpha-coronaviruses, including all SADS-CoV sequences, estimated using the RNA dependent RNA polymerase (RdRp) gene using and the GTR+F+I+Γ_4_ model of nucleotide substitution. The SADS-CoV node is collapsed and displayed in panel b. The host species of each virus is shown, and branches are colored by subgenera. Trees were midpoint rooted and bootstrap values >70% from 1000 bootstrap replicates are shown. (B) Maximum likelihood phylogeny (IQ-TREE) of SADS-CoV based on the whole viral genome utilizing the TIM2+F+I model and 1000 bootstrap replicates. SADS-CoV/Guangxi/2021 is denoted with a red pentacle. The tree is midpoint rooted and bootstrap values >70% are shown. (C) Reported outbreak geographical locations in China. Region in red with the pentacle (Guangxi) denotes the geographic origin of SADS-CoV/Guangxi/2021. The region in orange (Guangdong) represents the district of initial outbreak of SADS-CoV, while the regions in blue (Fujian) and green (Jiangxi) represent the locations of subsequent epidemics.

To address the issue of recombination in more detail we performed a variety of phylogenetic tests. Using the RDP4 package we identified one putative recombination event involving SADS-CoV/Guangxi/2021 that separated the ORF1ab from the spike genomic regions (*p*<0.05) (Figure 2). A subsequent analysis using two ML phylogenies estimated for the genome regions either side of the likely breakpoints revealed marked phylogenetic incongruence, indicative of potential recombination between ORF1ab and the rest of the viral genome. Specifically, phylogenetic analysis of the genomic region spanning nucleotides 1-20199 (i.e., the majority of the viral genome) revealed that SADS-CoV/Guangxi/2021 was most closely related, with 100% bootstrap support, to a sequence identified in Fujian province. In contrast, in a phylogeny of nucleotides 20200-27134 SADS-CoV/Guangxi/2021 clustered (with 74% bootstrap support) with sequences sampled from Jiangxi province (although the Fujian sequence is not available for this region) (Figure 2B).

**Figure 2.**
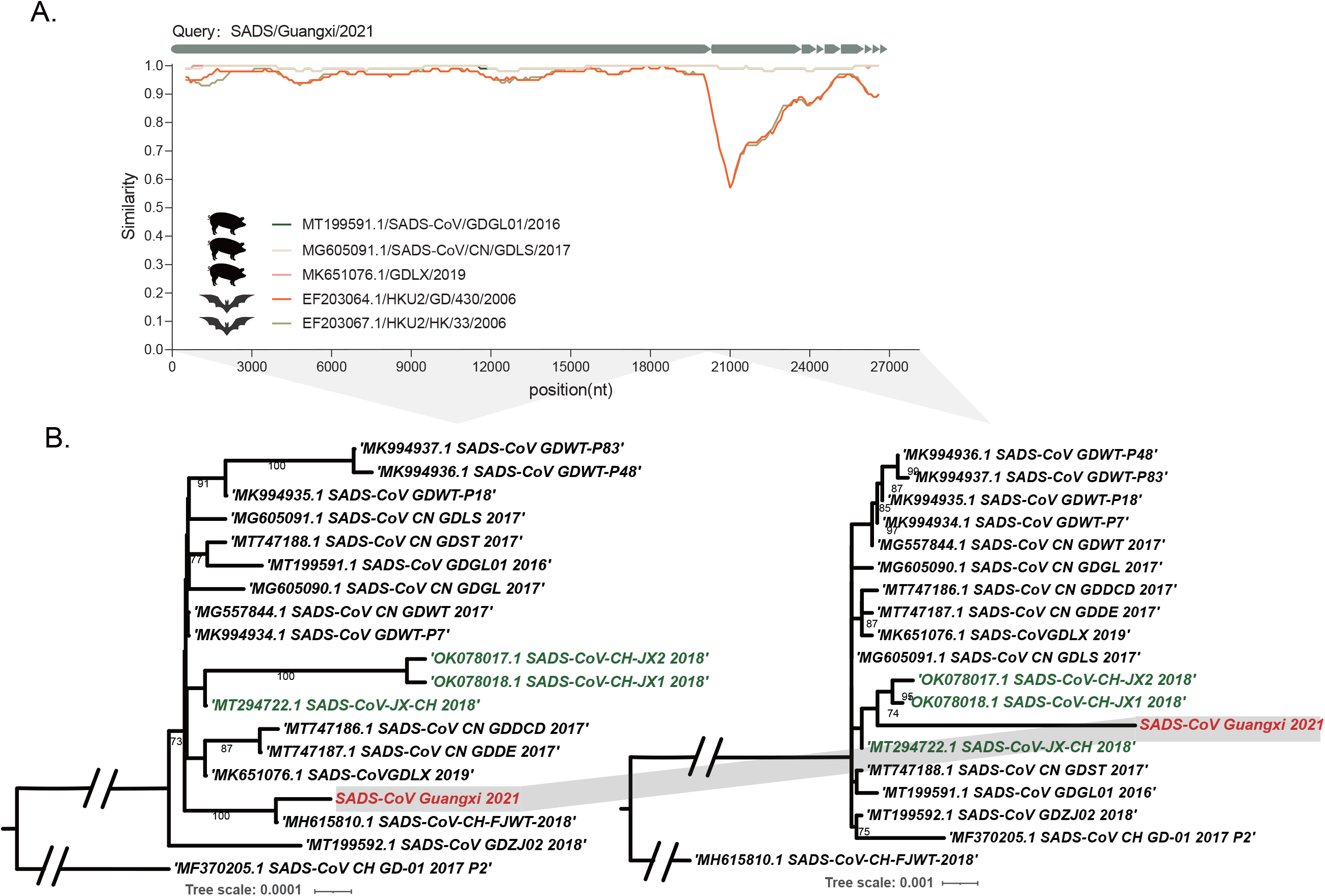
Recombination in SADS-CoV. (A) Genome-scale similarity comparisons of a query sequence (SADS-CoV/Guangxi/2021) against reference SADS-CoV sequences – SADS-CoV/GDGL01/2016 (green), SADS-CoV/CN/GDLS/2017 (yellow), and SADS-CoV/GDLX/2019 (pink) – as well as the bat-borne HKU2 coronavirus HKU2/HK/33/2006 (brown) and HKU2/GD/430/2006 (orange). This analysis used a window size of 1000 bp and a step size of 200 bp. (B) Maximum likelihood trees of either side of the likely recombination breakpoints (left: 1-20199 bp, right: 20200-27134). In the left tree, the best-fit substitution model was TIM+F+I, while in the tree on the right the equivalent model was TIM2+F+Γ_4_. Bootstrap values (>70%) are shown from 1000 bootstrap replicates, with grey shadow marking the topological incongruence.

## Conclusions

Using a combination of RT-qPCR and metagenomics we identified the re-emergence of SADS-CoV in Guangxi province, China, in infected pigs displaying severe and acute diarrhea and vomiting, weight loss, with a fatality rate up to 100% in piglets. Phylogenetic analysis suggested undetected circulation since at least 2019 as well as the inter-provincial spread of the virus. That the virus detected in Guangxi also has a recombinant origin further suggests that there is a diverse pool of SADS-CoVs in circulation that have yet to be identified.

No previous epidemics of SADS-CoV have been identified in Guangxi province (10). Strikingly, rather than disappearing, our surveillance and phylogenetic analysis suggests that this virus may have been circulating undetected since 2019, such that additional work should be directed toward finding the reservoir population(s). These findings also constitute an appropriate alert for the potential of this virus to have a devastating impact on the swine industry, as well as the possibility of future cross-species events to humans working at the human-animal interface (13–15).

## ACKNOWLEDGEMENTS

We thank Xiaoqin Xu and Yuli Luo, South China Agricultural university, for their generous assistance in the sampling and routine monitoring of SADS-CoV This research was funded by the Key-Area Research and Development Program of Guangdong Province [grant number 2019B020211003], Start-up Research Project of Maoming Laboratory (2021TDQD002), and China Agriculture Research System of MOF and MARA (cars-35). ECH is funded by an Australian Research Council Australian Laureate Fellowship (FL170100022).

## ETHICS STATEMENT

All sampling procedures were approved by the Animal Ethics Committee of South China Agricultural University and conducted under the guidance of the South China Agricultural University Institutional Animal Care and Use Committee (SCAU-AEC-2022A010).

## SUPPLEMENTARY FIGURE LEGENDS

**Figure S1. Phylogeny of Spike gene.** Maximum likelihood (IQ-TREE) phylogeny of the spike (S) gene utilizing the HKY+F model of nucleotides substitution. The tree is midpoint rooted and bootstrap values >70% are shown.

