## Appendix for "Re-emergence of Severe Acute Diarrhea Syndrome Coronavirus (SADS-CoV) in Guangxi, China, 2021"

**Methods	2**

**Supplementary Figure and Tables	4**

Figure S1	4

Table S1	5

Table S2	5

### Methods

**Sequence and Real Time-qPCR**

From winter 2019 to February 2021, we had launched a routine surveillance project of SADS-CoV in China, and we had sampled from the pig farms characterized by outbreaks of large-scale diarrhea in pigs. Totally, 1365 diarrhea feces and intestinal samples from twenty-three provinces, 5 autonomous regions, and 4 municipalities had been included in this study to perform real-time quantitative reverse transcription PCR (real-time qRT-PCR) to detect SADS-CoV, and there are no positive cases have been detected. Samples were kept in RNA keeper and transported to the laboratory in a dry ice environment. Then we stored it at -80℃ until processing. All the samples were further ground in were ground by a freezing grinder (JXFSTPRP-CLN-48, Shanghai Jingxin Industrial Development Co., Ltd., China) and RNA was extracted from the tissues using the RNAfast200 kit (Fastagen, Shanghai, China) according to the manufacturer’s instructions, and NGS was performed using a paired-end MGISEQ-200RS. We trimmed the adapter sequences of the short reads using Trimmomatic and removed low-quality reads (quality scores < 20), and then assembled the reads by MEGAHIT to obtain high-quality contigs. After assembly, the contigs were extracted and annotated to generate the whole genome, and then checked with BLAST, DIAMOND, and KRAKEN2. The consensus sequences of the whole genome were confirmed using Reverse Transcription-Polymerase Chain Reaction (RT-PCR) and Sanger sequencing. Amplification reactions were performed in a real-time PCR CFX96 system (Bio-Rad Laboratories, Hercules, CA, USA). Positive and negative controls and no template samples were included at random in each run. Data analysis and interpretation of results were performed with Seegene Viewer software allowed automated analysis and interpretation of results.

**Dataset**

All representative *Alaphcoronavirus* sequences and SADS-CoV sequences were downloaded from NCBI database. We further aligned the sequences using MAFFT v7.4, with deleting ambiguous regions using trimAl.

We have chosen five reference strains to make a similarity analysis using SimPlot, which showed that SADS-CoV/Guangxi/2021 shares high homogeneity with reference SADS-CoV genomes except for NS3a region. Also, it shared a high similarity among the ORF1ab region and the substantially heterogeneous S region with bat-borne HKU2.

**Recombination detection**

We constructed all the whole-genome sequences of SADS-CoV and HKU2 from the genebank with the novel isolate to do a full exploratory recombination scan using all methods mentioned in the phylogeny recombination. For our analysis, the following methods were considered: RDP, MAXCHI, CHIMAERA, 3SEQ, and GENECONV. We ran two additional approaches: BootScan and SiScan, as secondary methods. We used the recommended settings for all methods, and sequences that were classified as positive results by three or more methods showed *p*<0.05. The beginning and end of breakpoints identified with RDP4 were used to split the genome into regions for further phylogenetic analysis.

**Phylogeny analysis**

Phylogenetic analysis was constructed using IQ-TREE, with best-fit model of nucleotides substitution chosen according to Bayesian Information Criterion, with ultrafast bootstrap approximation with 1000 replicates.

**Supplement Figure 1.** **Phylogeny of Spike gene.**

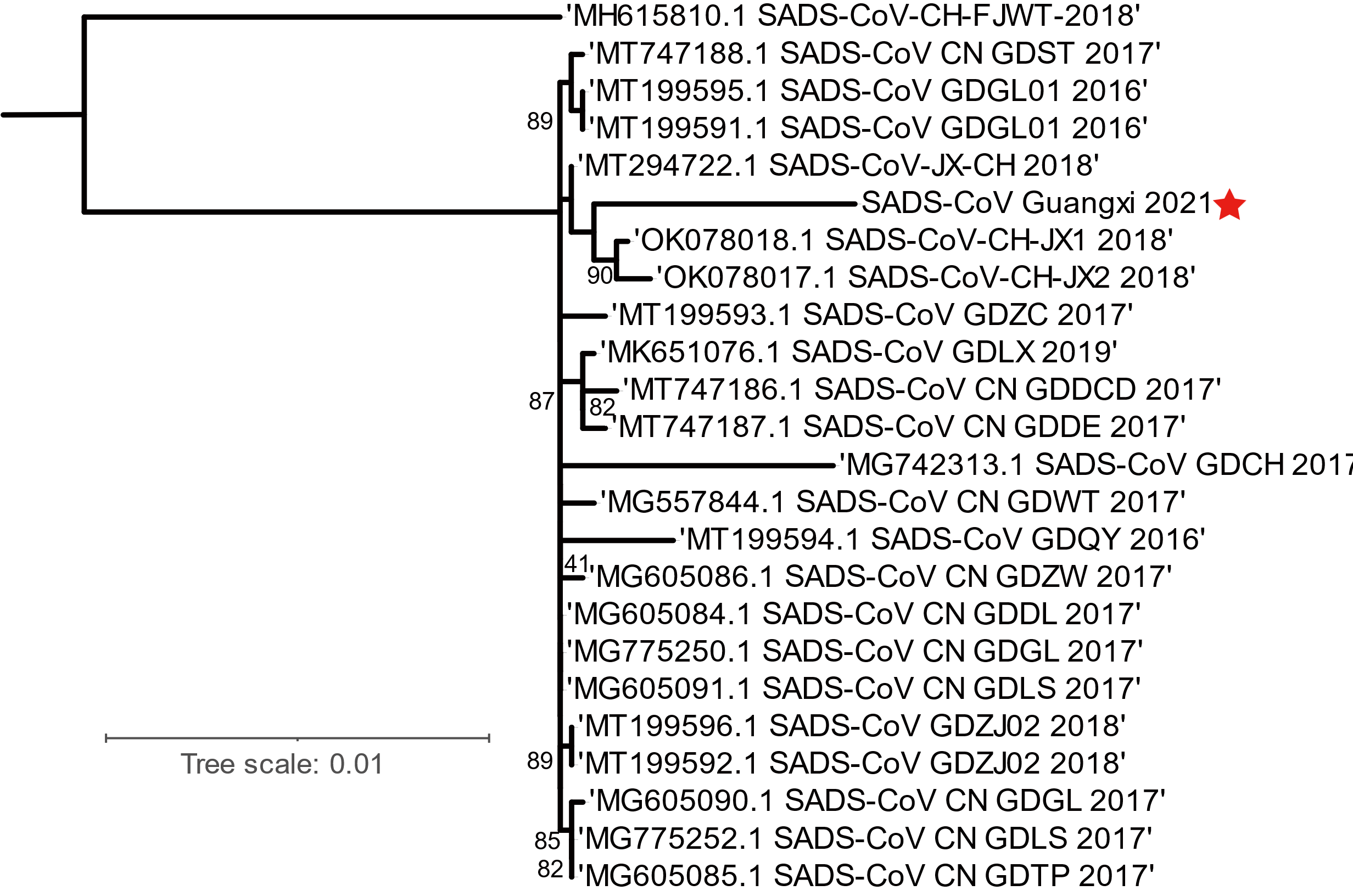

**Supplementary Table 1. Breakpoints of recombination.**

| Strain name | Regions | | | | | |  | Av. P-Val | | | | | | | |
| --- | --- | --- | --- | --- | --- | --- | --- | --- | --- | --- | --- | --- | --- | --- | --- |
|  | Major parent | Minor parent | Beginning  Breakpoint | 99% CI | Ending Breakpoint | 99% CI |  | | RDP | GENECONV | BootScan | MaxChi | Chimaera | SiScan | 3Seq |
| SADS-CoV/Guangxi/2021 | MT199592.1 | MK994935.1 | 20972 | 17326-23542 | 27132 | 26512-523 |  | | 3.246×10^-4^ | - | 9.332×10^-8^ | 5.378×10^-4^ | 1.281×10^-3^ | 6.374×10^-9^ | - |

**Supplementary Table 2. Characteristics of SADS-CoV identified in this study.**

| Library | Strain | Date | Region | CT | Accession number |
| --- | --- | --- | --- | --- | --- |
| 22R32984 | SADS-CoV/Guangxi/2021.5 | 2021.5 | Guangxi | 23.18 | ON911569 |
